# Disrupted Ecology and *H. parainfluenzae* Distinguish the Gut Microbiota of an Ethnic Minority Predisposed to Type 2 Diabetes Mellitus

**DOI:** 10.1101/2023.10.01.560393

**Authors:** Eric I. Nayman, Brooke A. Schwartz, Michaela Polmann, Alayna C. Gumabong, Max Nieuwdorp, Trevor Cickovski, Kalai Mathee

## Abstract

**Purpose:** Decreased gut microbiota production of short-chain fatty acids (SCFAs) has been implicated in type 2 diabetes mellitus (T2DM) disease progression. Most microbiome studies focus on ethnic majorities. This study aims to understand microbiome differences between an ethnic majority (the Dutch) and minority (the South-Asian Surinamese (SAS)) group with a lower and higher prevalence of T2DM, respectively.

**Methods:** Microbiome data from the Healthy Life in an Urban Setting (HELIUS) cohort were used. The 16S rRNA V4 region was sequenced. Two age– and gender-matched groups were compared: the Dutch (n = 41) and SAS (n = 43). Microbial compositions were generated via DADA2. Alpha and beta-diversity and Principal Coordinate Analysis (PCoA) were computed. DESeq2 differential bacterial abundance and LEfSe biomarker analyses were performed to determine discriminating features. Co-occurrence networks were constructed to examine gut ecology.

**Results:** A tight cluster of bacterial abundances was observed in the Dutch women, which overlapped with some of the SAS microbiomes. The Dutch gut contained a more interconnected microbial ecology, whereas the SAS network was dispersed. *Bacteroides caccae, Butyricicoccus, Alistipes putredinis, Coprococcus comes*, *Odoribacter splanchnicus,* and *Lachnospira* characterized the Dutch gut. *Haemophilus*, *Bifidobacterium,* and *Anaerostipes hadrus* characterized the SAS gut. All but *Lachnospira* and certain strains of *Haemophilus* are known SCFA producers.

**Conclusion:** The Dutch gut microbiome was distinguished from the SAS by diverse, differentially abundant SCFA-producing taxa with significant cooperation. The dynamic ecology observed in the Dutch was lost in the SAS. The higher prevalence of T2DM in the SAS may be associated with the dysbiosis observed.

## Background

A microbiome is a community of phylogenetically diverse microorganisms and their multi-omic content that inhabit a specific ecological niche, such as the human gut [2]. The microbiome and the host are embedded in a mutually dependent relationship that impacts host behavior and microbial community structure and function [3]. As host-microbe interactions shape reciprocal fitness, phenotype, and metabolism, the host and the microbiome coevolve [2,3]. *Dysbiosis*, or pathologic alteration of the baseline microbial milieu, may drive and/or result from disease progression [4,5]. Our resident microbial flora has been widely recognized as a key but not yet fully understood mediator in the pathophysiology of many communicable and noncommunicable diseases [6–12], including type 2 diabetes mellitus (T2DM) [13–17].

Diabetes is a chronic, metabolic disease in which hyperglycemia leads to multi-organ damage over time [18]. The disease burden of T2DM has been increasing globally. Since the 1990s, the prevalence of T2DM has increased, and the age-adjusted prevalence is expected to rise from 6.3% in 2019 to 7.8% in 2045 across Europe [19]. Disproportionately high rates of diabetes and related complications affect migrant and ethnic minority groups living in Western societies [20]. Diet, genetic predisposition, body weight, and sedentary lifestyle are key factors in the multifaceted pathophysiology of T2DM [21]. Recently, the gut microbiome of diabetics has been shown to be distinctly different from that of normoglycemic, insulin-sensitive individuals [15–17,22]. Most agree that the abundances of *Ruminococcus, Fusobacterium*, and *Blautia* are positively correlated with T2DM while *Bifidobacterium, Bacteroides, Faecalibacterium, Akkermansia,* and *Roseburia* are negatively correlated with T2DM [14,23–25]. However, a large knowledge gap still exists: how do the quantitative presences of the many other gut inhabitants vary with hyperglycemia and insulin resistance [24]? For example, many disagree on the nature of the correlative relationship between T2DM and the abundance of *Lactobacillus* [24].

Short-chain fatty acids (SCFAs), namely, acetate, butyrate, and propionate, and the microbes that produce them have been of particular interest in the diabetic gut because of the favorable effect that these molecules have on host function [26–29]. Most of the beneficial effects of SCFAs on glucose metabolism and insulin signaling are mediated via the GPR41 and GPR43 receptors [28,30,31]. These molecules can activate intestinal gluconeogenesis [28,32], potentiate glucose-stimulated insulin secretion through glucagon like peptide-1 (GLP-1) dependent and independent pathways [30,31,33], and attenuate the chronic release of pro-inflammatory cytokines that worsens insulin resistance [34–36]. SCFAs have many other important functions on host metabolism, which are well described by recent reviews [27,28]. Ultimately, a decrease in SCFA-producing taxa is comorbid with T2DM and may cause or worsen the disease. Of note, cross-feeding mechanisms between SCFA-producing taxa play a major role in the gut microbial functional ecology. For example, *Bacteroides thetaiotaomicron, Blautia obeum, Roseburia inulinivorans, Listeria* sp., and *Clostridium sphenoides* can all create a major intermediate metabolite (1,2-propanediol) which can be used by *Lactobacillus reuteri* to make propionate. Additionally, *Roseburia* can take up acetate, the most widely produced SCFA, and make butyrate from glucose [37].

Ethnicity and place of habitation are thought to play an even larger role than metabolic health in shaping gut microbiome composition [38–40]. Ethnicity is a particularly important factor as it connotes similar diet, shared genetics, and migration patterns, all of which have their own variable impact on gut microbial flora. The multi-ethnic Healthy Life in an Urban Setting (HELIUS) prospective cohort study estimated that the contribution of ethnicity on gut microbiome composition is ∼6% [39]. The HELIUS cohort included participants from the six major ethnic groups living in Amsterdam, the Netherlands at the time of sample collection. Of these ethnicities, the Dutch, Ghanaian, and South-Asian Surinamese (SAS) were found to have the most discriminant gut microbiomes [39]. In the Dutch population, the ethnic majority of Amsterdam, nine core gut bacterial species were identified: *Subdoligranulum* sp., *Alistipes onderdonkii*, *Alistipes putredinis*, *Alistipes shahii*, *Bacteroides uniformis*, *Bacteroides vulgatus*, *Eubacterium rectale*, *Faecalibacterium prausnitzii* and *Oscillibacter* sp. These were also consistently found across many other populations [40]. Of these, several were shown to be significantly depleted, e.g., *Faecalibacterium prausnitzii,* and enriched, e.g., *Bacteroides* sp., in the SAS as compared to the Dutch [39].

The aim of our study is to compare the gut microbiome ecology between the Dutch and SAS groups, as these had the lowest and highest prevalence of T2DM among the HELIUS ethnicities, respectively. Our objective is to characterize and replicate, as microbiome studies are notoriously challenged with reproducibility, the differential bacterial biomarkers and inter-taxa relationships between these two groups. We demonstrate differences in the SCFA-producing taxa and correlate these with ethnicity and metabolic health.

## Methods

### Genomic Data Source

Our dataset is from a multi-ethnic prospective cohort study, the HELIUS study. This cohort is composed of four major ethnic groups (Surinamese, Dutch, Ghanaian, Moroccan, and Turkish) aged 18 to 70 years old (Table 1). Participants were randomly invited, and then stratified by ethnicity. All were living in Amsterdam, the Netherlands at the time of sample collection (2011 – 2015). A total of 2,170 stool samples, each from a different individual, were collected, and the metagenomes were sent for sequencing of the 16S rRNA V4 hypervariable region. This was done on a 2 x 250 bp MiSeq system with use of the 515F and 806R primers. The 2,170 sequenced fecal samples yielded a total raw read count of 177,089,775. Further detail about cohort composition, data collection, and gene sequencing protocols have been previously described [39,41]. The HELIUS study was approved by the medical ethics committee of the Amsterdam University Medical Center, and all participants provided informed consent prior to enrollment in the study.

**Table 1.** HELIUS Microbiome Cohort.

**(a) Six Major Demographics Represented in the HELIUS Microbiome Dataset**
| Ethnicity | Male | Age Range (years old) | Female | Age Range (years old) |
| --- | --- | --- | --- | --- |
| Dutch | 229 | 22 - 71 | 211 | 19 - 71 |
| Turkish | 116 | 19 - 69 | 83 | 19 - 68 |
| African Surinamese | 173 | 27 - 71 | 272 | 22 - 70 |
| South-Asian Surinamese | 162 | 26 - 70 | 196 | 18 - 71 |
| Ghanaian | 187 | 21 - 68 | 181 | 19 - 65 |
| Moroccan | 156 | 19 - 69 | 126 | 19 - 68 |
| Total | 1023 | 19 - 71 | 1069 | 18 - 71 |

**(b) Sub-Cohort Used in this Study**
| Ethnicity | Male | Female | Total | Age Range (years old) |
| --- | --- | --- | --- | --- |
| Dutch | 17 | 24 | 41 | 53 - 55 |
| South-Asian Surinamese | 21 | 22 | 43 | 53 - 55 |
| Total | 38 | 46 | 84 | 53 - 55 |

### Ethnicity

A person was defined as of non-Dutch ethnic origin if he/she fulfilled one of two criteria: (1) he/she was born outside of the Netherlands and had at least one parent born outside the Netherlands (first generation) or (2) he/she was born in the Netherlands but both parents were born outside of the Netherlands (second generation). For the Dutch samples, people who were born in the Netherlands and whose parents were born in the Netherlands were invited. The country of birth indicator for ethnicity was limited in that people who were born in the same country might be of different ethnic background, which in the Dutch context was applicable to the Surinamese population. Therefore, after data collection, participants of Surinamese ethnic origin were further classified by self-reported ethnic origin (obtained by questionnaire) as ‘African’, ‘South-Asian’, ‘Javanese’, or ‘other’.

### Microbial Community Composition

Paired-end reads were input into a DADA2 (v1.16) [42] workflow to generate microbial compositions. All parameters were kept at default values except those described here. First, primers were removed from sequence reads. Then, reads were quality filtered and truncated. Two base-pair (bp) errors were allowed per 250 bp read, and read-ends were trimmed down to a quality score threshold of 30. Then, reads were dereplicated and merged. A minimum overlap of 30 bp was required for merging to occur. After merging, chimeric sequences were removed from the amplicon sequence variants (ASVs). Lastly, taxonomy was assigned via queries to the SILVA 16S rRNA gene reference database (v138.1) [43]. A paired, two-tailed t-test was calculated between the relative abundances (RAs) of each taxon to determine if significant differences in a particular taxon existed between the two ethnic groups.

### Principal Coordinate Analysis

To estimate the degree of differentiation between the common core microbiota [44] of the Dutch and SAS samples, Principal Coordinate Analysis (PCoA) [45] was applied based on the RAs of the taxa. First, a prevalence threshold of 50% [46] was applied. Then, the first two principal components were plotted in a two-dimensional space.

### Microbiota Diversity

To measure the microbial diversity within each sample, the following alpha diversity indices were computed: Chao richness [47], Shannon [48], Fisher [49], and inverse Simpson [50]. Weighted UniFrac distances [51,52] were computed to estimate beta diversity, or how different the samples within each ethnic group were from one another in terms of phylogeny and abundance.

### Biomarker Analyses

To identify potential biomarkers that could distinguish the Dutch and SAS microbiota, a linear discriminant analysis of effect size, or LEfSe analysis [53], was performed (*P*<0.05 and LDA effect size>1). LEfSe proposes microbial biomarkers based on relative abundance, effect size, and biological consistency. It can also rank the significance of the biomarkers, i.e., taxa, because it calculates the effect size of each. This rank of significance is provided via the LDA score. LEfSe is especially useful for determining significant differences in taxa with low RAs, which is difficult to do using pure abundance data alone.

Additionally, a differential expression analysis for sequence count data, or DESeq2 [54,55], algorithm was used to calculate if significant differences existed between the bacterial abundances of the two ethnic groups. Lastly, to determine likely species– and strain-level taxonomic identities of the proposed biomarkers, the nucleotide sequences of the corresponding ASVs were input into a BLAST [56] search for similar sequences contained only in rRNA/ITS databases.

### Network Analysis

To estimate two-way ecological relationships in the Dutch and SAS microbiota [57], we built microbial co-occurrence (social) networks [58] using microbial RA data at the lowest possible taxonomic classification level. We computed SparCC [59] correlations between each pair of taxa (*P*<0.01). Results were displayed as a network, with nodes representing taxa (size proportional to RA), and edges representing correlation (green=positive, estimating cooperation; red=negative, estimating competition). Networks were visualized using the Fruchterman-Reingold algorithm [60] to clarify community structure.

## Results

### Cohort Analysis

This study analyzed a strictly age– (53 to 55 years-old) and gender-matched subset of the HELIUS gut microbiome cohort (Table 1) to account for any potential confounding effects of these variables on gut microbiota [61,62]. Only the Dutch and SAS ethnicities were analyzed because these were reported to have the lowest (five %) and highest (21.5%) prevalence of T2DM, respectively, among the ethnic groups of the HELIUS study [63]. After matching and adjusting for the high computational demand of bioinformatic processing, 41 Dutch and 43 SAS subjects were analyzed.

### Diversity Analyses

To estimate the degree of differentiation between the common core microbiota of each ethnicity, PCoA [45] was applied based on the RAs of the top 25 most abundant taxa (Fig. 1a). The Dutch had more similar core gut microbiota to each other as far as abundance goes. Contrarily, the SAS varied much more from person to person. Qualitatively, the SAS abundances formed two distinct clusters while the Dutch formed a single, tight cluster. Within the Dutch cluster, the Dutch females formed an even tighter cluster. There was some overlap between both ethnicities and genders.

**Fig. 1.**
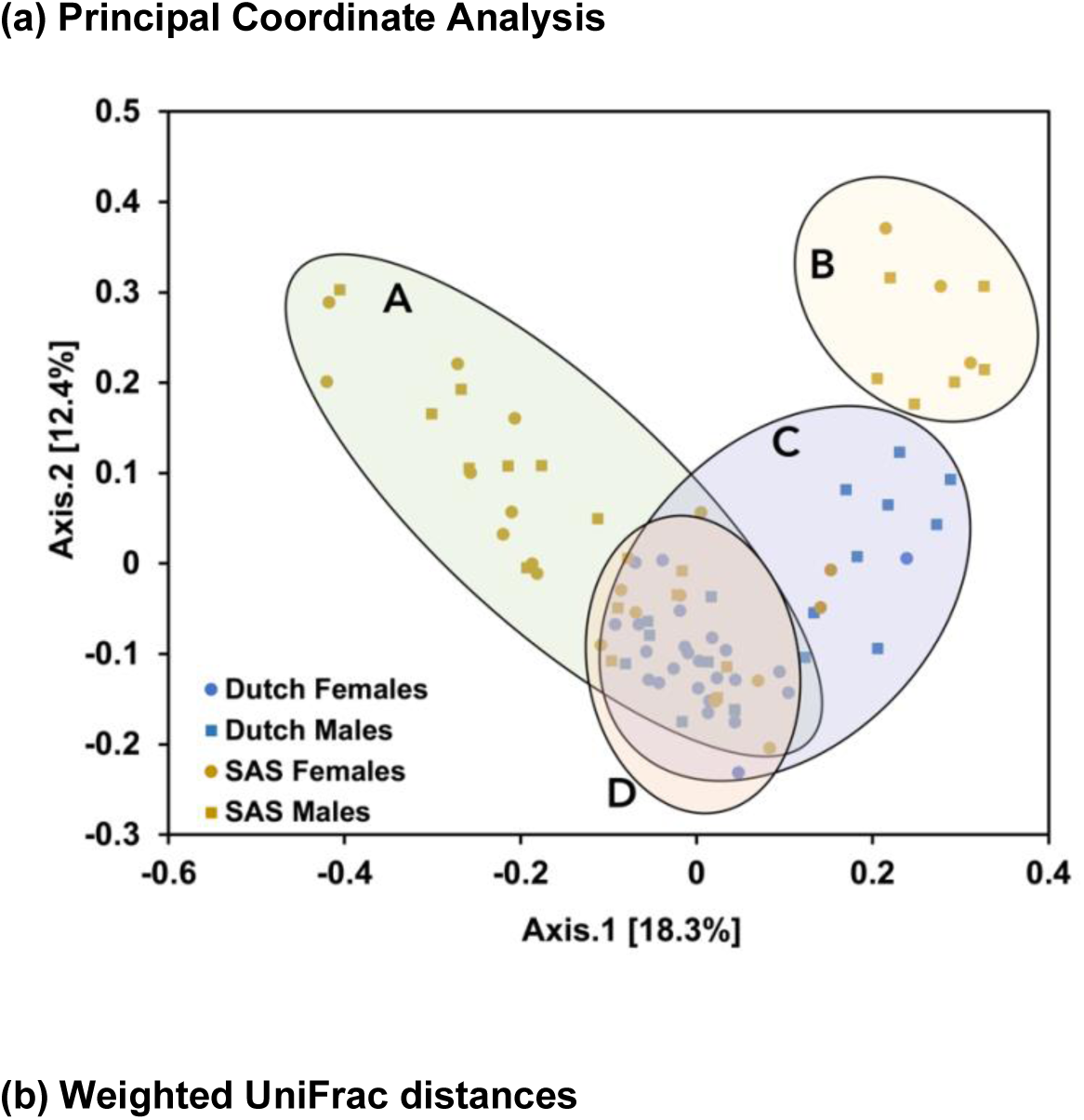
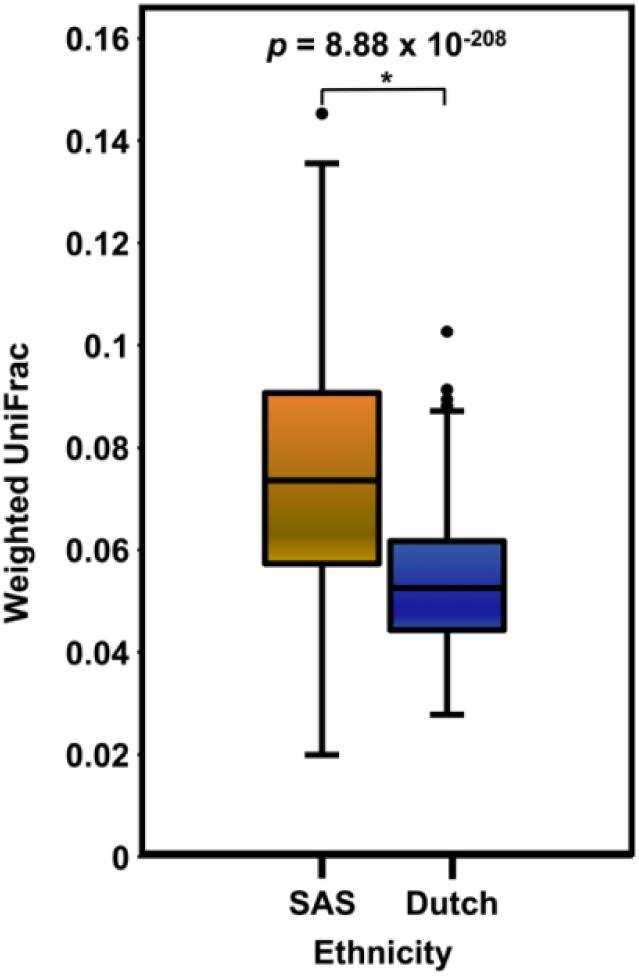
Microbial diversity between samples: (a) Principal Coordinate Analysis. To estimate the degree of differentiation between each ethnicity’s common core microbiota, which was approximated by the top 25 most abundant taxa, PCoA [45] was applied based on the relative abundances. First, a prevalence threshold of 50% [46] was applied. Then, the first two principal components were plotted in a two-dimensional space. The PCoA showed that the Dutch had more similar gut microbiomes regarding the quantitative presence of the common core taxa. Contrarily, the SAS varied much more from person to person. Qualitatively, the SAS formed two clusters (A&B), and the Dutch formed a single cluster (C). Microbiota from the Dutch women formed the tightest cluster (D). There is some overlap between all the groups. **(b) Weighted UniFrac distances** To estimate beta diversity, or how different the samples within each ethnic group were from one another for phylogeny and abundance, weighted UniFrac distances [51,52] were computed. On average, the Dutch had a significantly more consistent gut microbiota between samples than did the SAS (*P* = 8.88 x 10^-208^)

Four different metrics of alpha diversity were computed (Table 2) to achieve consensus on the overall trend in microbial diversity across the samples, as each measurement of alpha diversity is uniquely biased [64]. All metrics indicated that the Dutch gut harbored a significantly more diverse microbial milieu (*P*<0.001). Weighted UniFrac distances [51,52] between samples were computed to approximate beta diversity (Fig. 1b). A lower UniFrac distance indicates a greater degree of similarity between two sets of taxa in terms of phylogeny and abundance. The Dutch had significantly more similar gut bacterial communities to each other than did individuals of SAS origin (*P* = 8.88 x 10^-208^).

**Table 2.** Alpha diversity.

| Alpha Diversity Metric | South-Asian Surinamese | Dutch | *p*-value |
| --- | --- | --- | --- |
| Chao1 | 151.51 | 267.8 | $1 \times 10^{-14}$ |
| Shannon | 3.7 | 4.43 | $2.27 \times 10^{-13}$ |
| Fisher | 19.39 | 36.29 | $2.57 \times 10^{-14}$ |
| Inverse Simpson | 23.15 | 44 | $1.55 \times 10^{-10}$ |

### Microbial Abundances

DADA2 [42] was used to infer gut microbiome compositions. The common core gut microbiota of the Dutch and SAS were represented by the top 25 most abundant genera (Fig. 2), or otherwise lowest possible phylogenetic rank, because of the limitations of identifying finer phylogenetic resolutions with short-read, targeted 16S sequencing [44,65]. Several tests of differential abundance between the two ethnicities were performed because methods for approximating differential abundance largely vary in output. So, it is best to perform several to attain consensus on the differentially abundant taxa, or potential biomarkers [66].

**Fig. 2.**
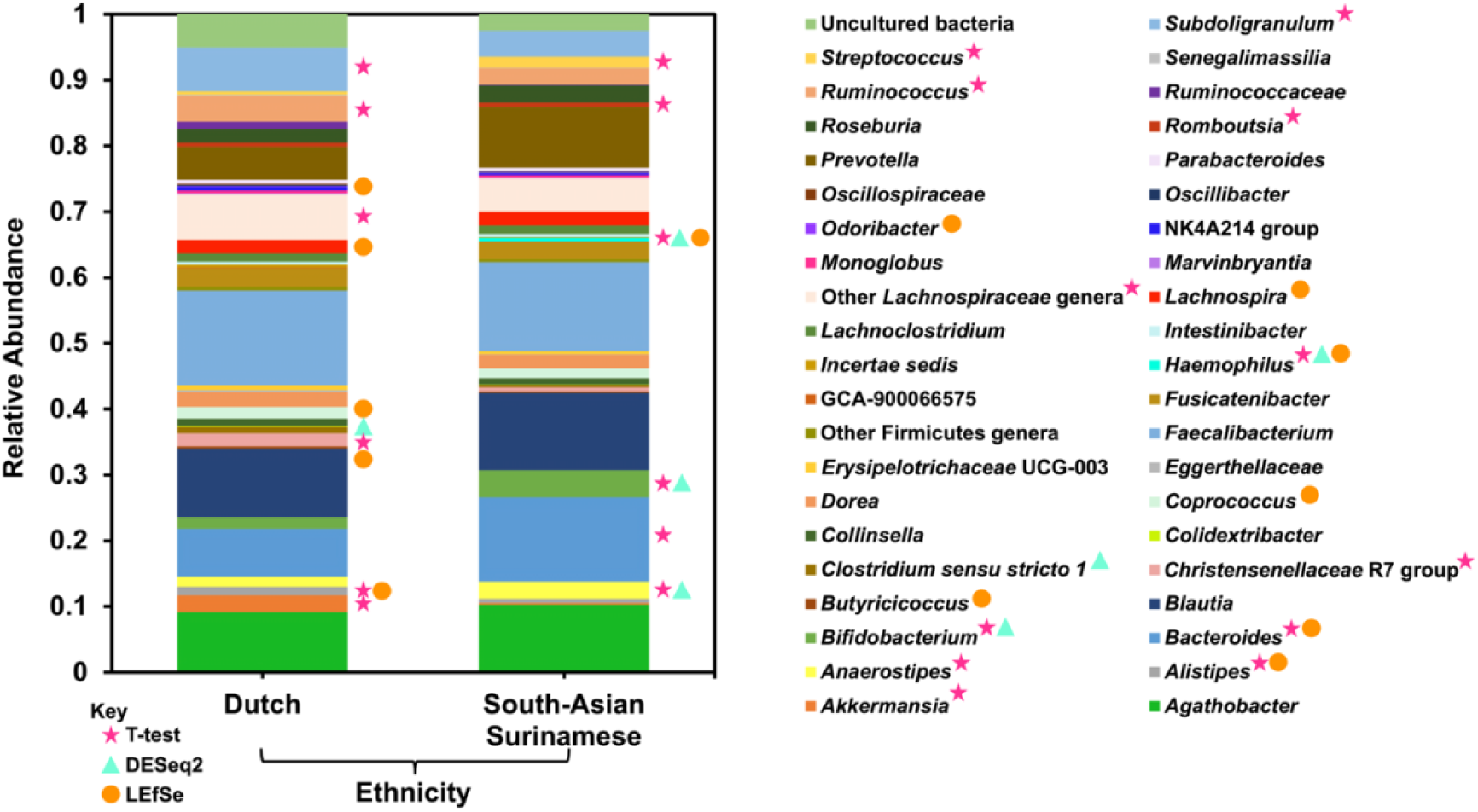
Relative abundances of common core gut microbiota: The common core gut microbiota of the Dutch and South-Asian Surinamese ethnic groups were represented by each ethnicity’s 25 most abundant genera, or otherwise lowest possible phylogenetic rank. Relative abundances were computed by DADA2 (v1.16) [42]. The pink stars denote the taxa detected as significantly differentially abundant by a DESeq2 [54,55] analysis, the blue triangles denote those identified as potential biomarkers by LEfSe, and the orange circles represent taxa found to be in significantly different abundance by a paired, two-tailed t-test. Taxa highlighted by more than one test of differential abundance are more likely to truly differentiate the two groups

First, a paired, two-tailed t-test between each of the shared common core taxa was calculated (*P*<0.05). The following taxa had significantly higher average RAs in the Dutch guts: *Subdoligranulum, Ruminococcus*, a group of unclassified *Lachnospiraceae* genera, *Christensenellaceae* R7 group, *Alistipes*, and *Akkermansia*. The following taxa had significantly higher average RAs in the SAS guts: *Streptococcus, Romboutsia, Haemophilus, Bifidobacterium, Bacteroides*, and *Anaerostipes*.

The second test of differential abundance performed was a DESeq2 analysis (Table 3). In the Dutch samples, *Clostridium sensu stricto 1* was found to be in significantly higher abundance while *Bifidobacterium, Haemophilus,* and *Anaerostipes* were found to be in significantly higher abundances in the SAS samples (*P*-values<0.05).

**Table 3.** DESeq2 Differentially Abundant Taxa.

| Dutch | South-Asian Surinamese |
| --- | --- |
| *Clostridium sensu stricto* 1* | *Bifidobacterium* * |
| *Akkermansia* | *Haemophilus* * |
| UCG-002 | *Anaerostipes* * |
| *Intestinibacter* | *Bacteroides* |
| *Eggerthellaceae* | *Christensenellaceae* R7 Group |
| *Erysipelotrichaceae* UCG-003 | *Blautia* |
| *Coprococcus* | *Lachnospira* |
| *Marvinbryantia* | *Oscillibacter* |
| *Firmicutes* | *Parabacteroides* |
| *Dorea* | *Lachnospiraceae* UCG-004 |
| *Oscillospiraceae* | *Monoglobus* |

Lastly, a LEfSe [53] biomarker analysis was computed (Fig. 3). The LDA scores assigned by the LEfSe estimate the individual contribution of each bacterium to the overall uniqueness of the community. Per the LEfSe, the discriminating bacterial biomarkers of the Dutch gut included *Odoribacter splanchnicus, Lachnospira, Bacteroides caccae, Alistipes putredinis, Coprococcus comes,* and *Butyricicoccus*. *Haemophilus* and its ascending phylogenetic ranks characterized the SAS gut.

**Fig. 3.**
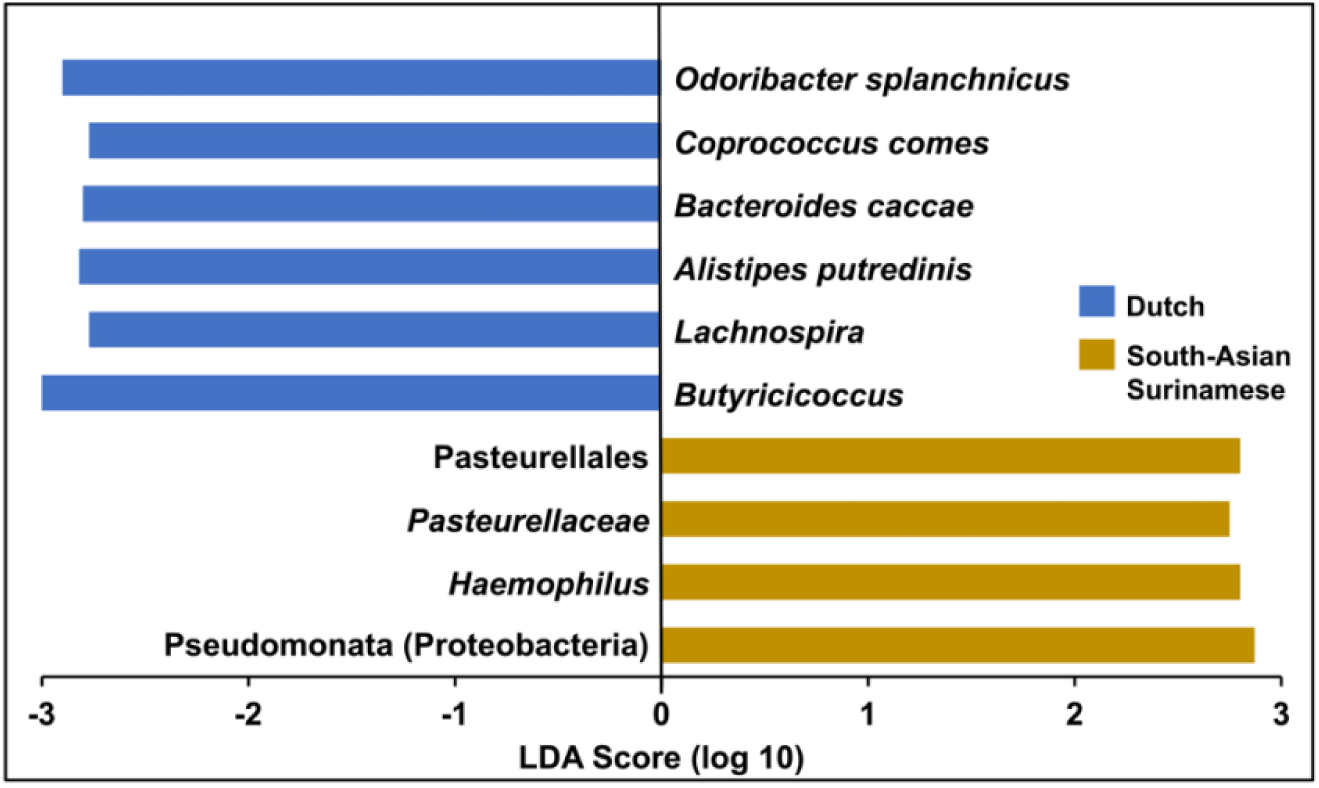
LEfSe biomarker analysis: To identify potential microbial biomarkers that could distinguish the Dutch and SAS gut microbiomes, a LEfSe analysis [53] was performed (*P*<0.05 and LDA effect size>1). The LEfSe was set to propose discriminating taxa at the lowest possible phylogenetic level. The LDA score for each potential biomarker is displayed as a histogram, with all scores falling between |2.5 – 3|. The proposed biomarkers of the Dutch gut microbiome included *Odoribacter splanchnicus, Coprococcus comes, Bacteroides caccae, Alistipes putredinis, Lachnospira,* and *Butyricicoccus*. *Haemophilus* and its ascending phylogenetic ranks discriminated the SAS gut

The full phylogenetic lineage of each differentiating taxon proposed by the DESeq2 and LEfSe analyses is provided (Fig. 4). We also inferred the most likely species– and strain-level identities of the differentially abundant taxa by performing a BLAST [56] search of the nucleotide sequence of each corresponding ASV uncovered in our sequencing (Table 4). There was notably more species and strain diversity in *Clostridium sensu stricto 1* and *Bifidobacterium* as compared to all the other discriminating taxa.

**Fig. 4.**
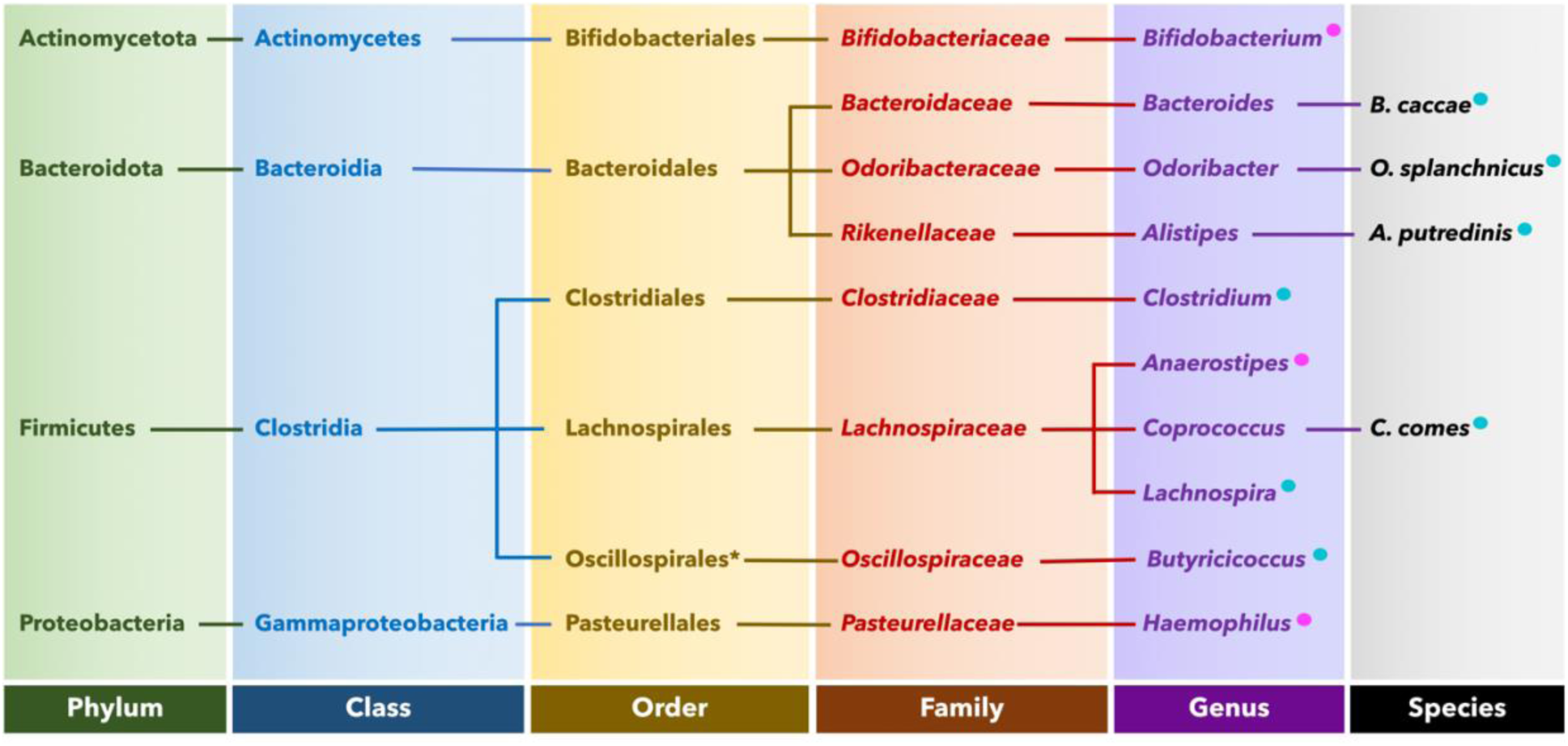
Phylogenetic tree of the proposed gut microbial biomarkers: The phylogenetic lineages of the taxa that were found to distinguish the gut microbiomes of the Dutch and SAS ethnic groups, as per the LEfSe [53] or DESeq2 [54,55] analyses, are shown. The blue circles represent microbes that are potential biomarkers of the metabolically healthier Dutch, and the pink circles represent those of the more T2DM-afflicted SAS

**Table 4.**
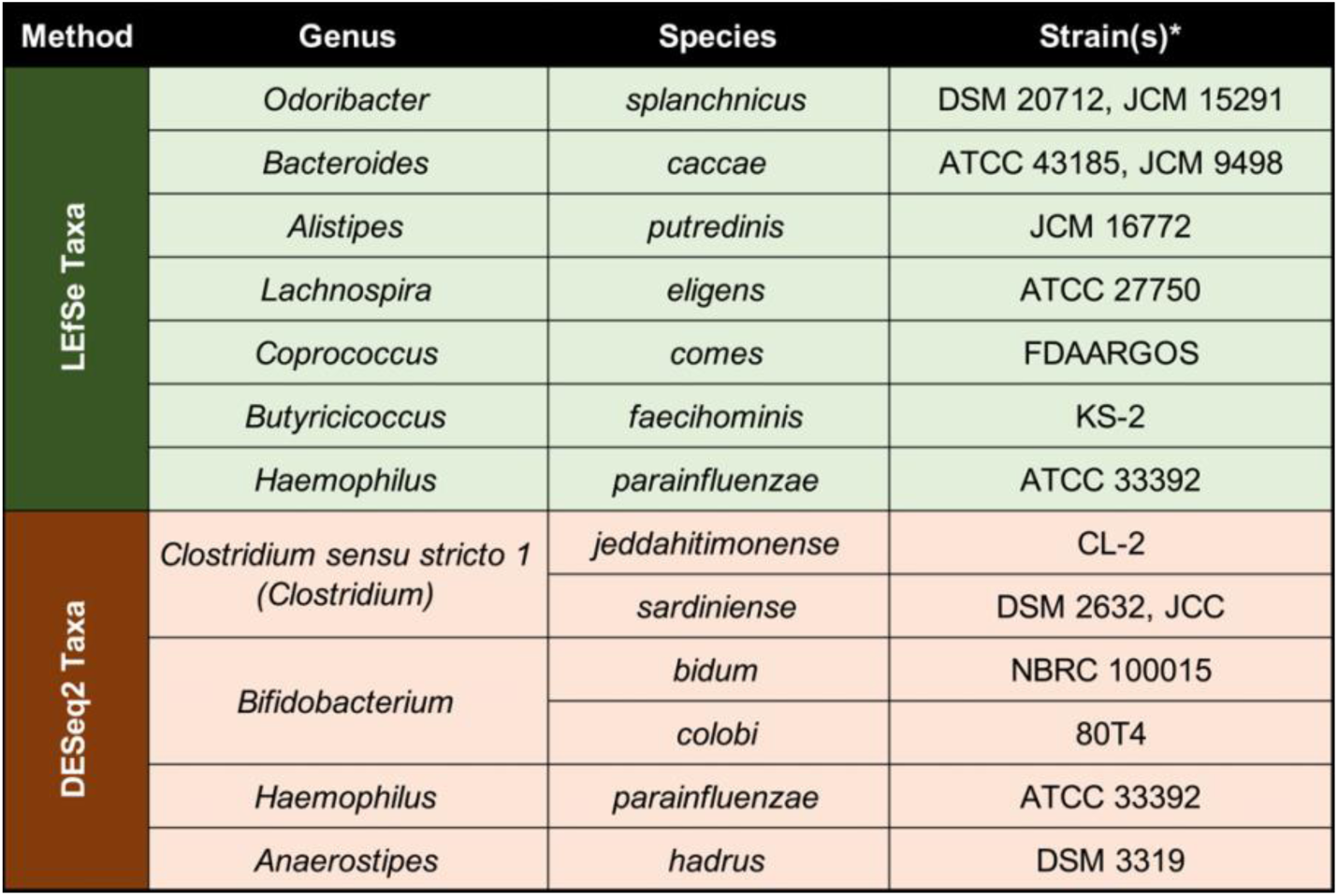
Species– and strain-level identities of the differentiating taxa.

The per sample distributions of the RAs of the differentially abundant taxa, as determined by either a paired, two-tailed t-test, DESeq2 analysis, and/or LEfSe biomarker analysis was then plotted (Fig. 5). *Clostridium sensu stricto 1* was entirely absent from the SAS guts while *Streptococcus* and *Romboustia* were totally missing from the Dutch guts. Some bacteria significantly varied in RA between individuals of one ethnicity. *Bacteroides* showed the most person-to-person variation in both ethnic groups, but this was more pronounced in the SAS. *Subdoligranulum* was widely distributed across the SAS. The RAs of *Streptococcus* in the SAS gut had the most outliers. The middle quartiles of the RAs for a group of unclassified *Lachnospiraceae* genera were almost completely non-overlapping between the two ethnic groups.

**Fig. 5.**
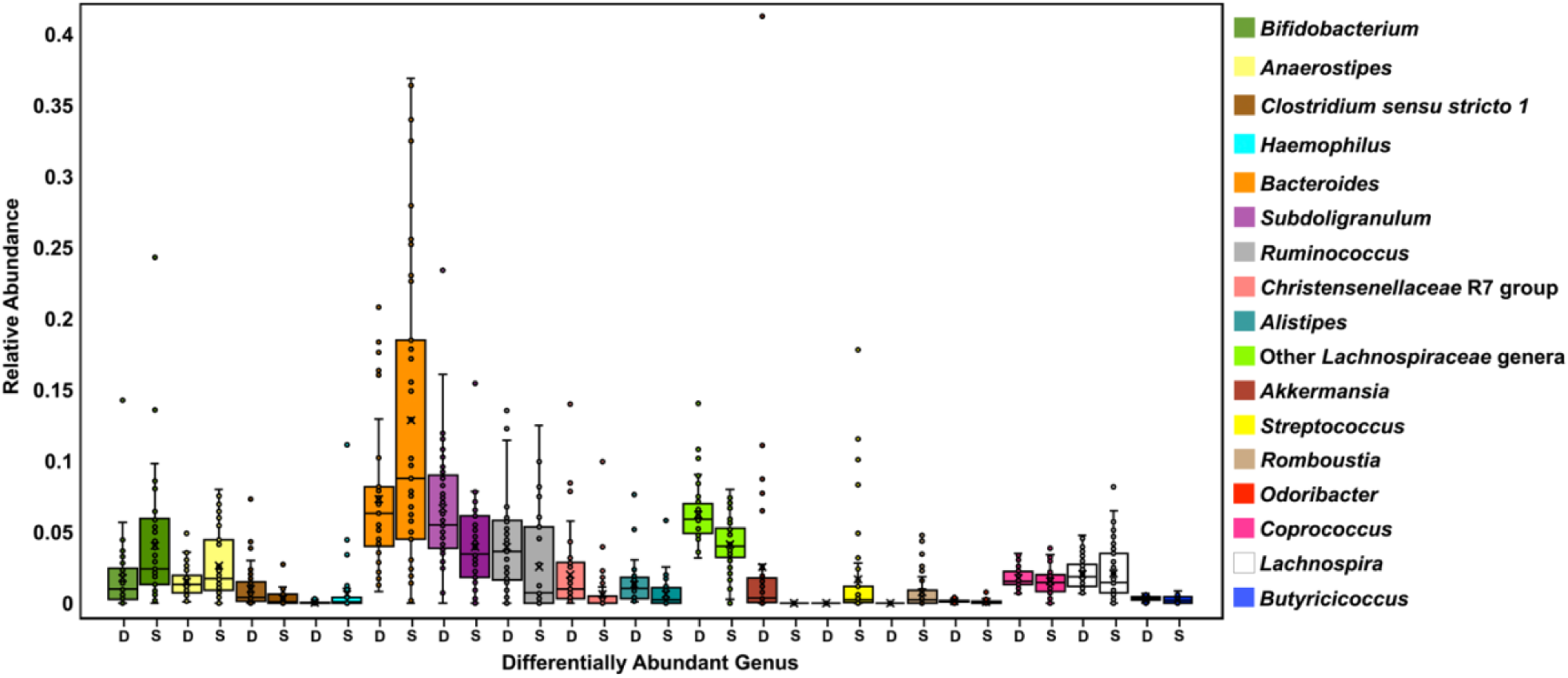
Per sample distribution of relative abundances of differentiating taxa: The per sample distribution of the RAs of the differentially abundant genera (*P*-values<0.05), as determined by either a paired, two-tailed t-test, DESeq2 analysis, and/or LEfSe biomarker analysis was plotted as a box and whisker plot. The t-test highlighted *Subdoligranulum, Ruminococcus*, a group of unclassified *Lachnospiraceae* genera, *Christensenellaceae* R7 group, *Alistipes*, and *Akkermansia* as having significantly RAs in the Dutch while *Streptococcus, Romboutsia, Haemophilus, Bifidobacterium, Bacteroides*, and *Anaerostipes* as having significantly higher RAs in the SAS gut. The DESeq2 analysis found *Clostridium sensu stricto 1* as differentially abundant in the Dutch and *Bifidobacterium, Haemophilus,* and *Anaerostipes* as differentially abundant in the SAS samples. Per the LEfSe, the discriminating biomarkers of the Dutch gut included *Odoribacter splanchnicus, Lachnospira, Bacteroides caccae, Alistipes putredinis, Coprococcus comes,* and *Butyricicoccus* while *Haemophilus* characterized the SAS gut

### Co-occurrence Network Analyses

To depict the ecological framework of each ethnic group’s gut microbiome, bacterial co-occurrence networks were constructed (Fig. 6), as previously described [58]. The thickness of the lines, or edges, is indicative of correlative strength. Green lines represent positive and red lines represent negative correlations. The number next to some of the organisms is a centrality score [67], which indicates how important that taxon is to the overall community structure. At a macroscopic view, the community is tightly knit with many positive correlational relationships between taxa in the Dutch. This interconnectedness is lost in the SAS.

**Fig. 6.**
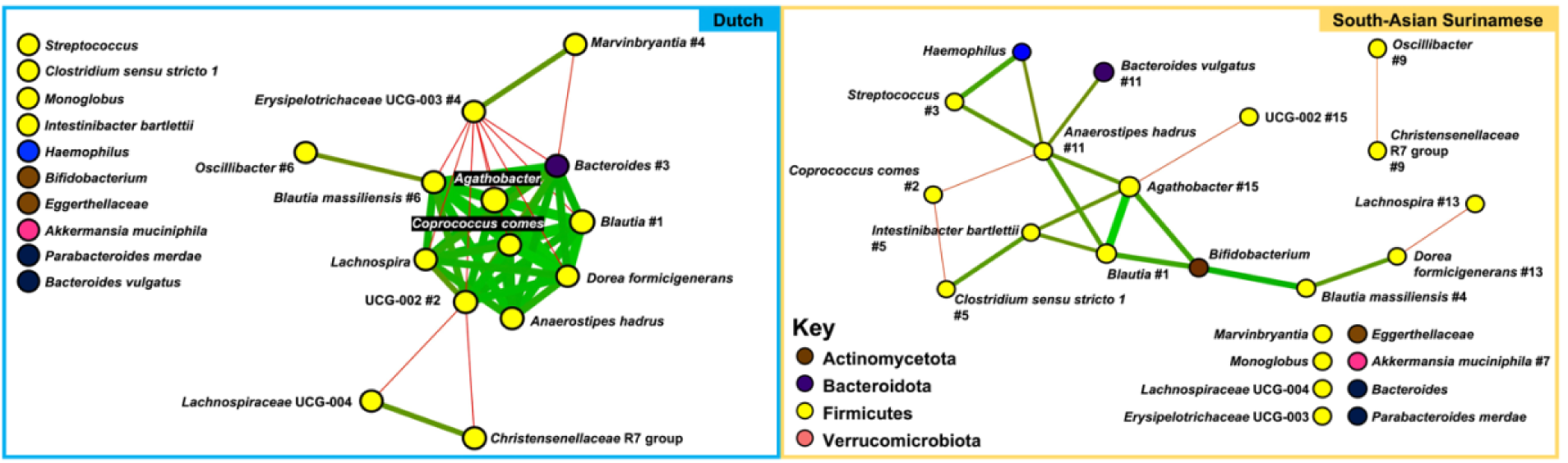
Bacterial co-occurrence networks: Bacterial co-occurrence (social) networks of the Dutch and SAS gut microbiomes are shown. RA data was used at the species or next lowest possible taxonomic classification level. These networks were constructed from pairwise correlation (SparCC [59], *P*<0.01) matrices using Cytoscape, and visualized using the Fruchterman-Reingold algorithm [60] to clarify community structure. Each node is a distinct ASV, and the node colors represent different phyla, as shown in the key. The thickness of the lines, or edges, indicates correlative strength, with green lines representing positive and red lines representing negative correlations. The number next to many organisms is a centrality score, which indicates how important that taxon is to the overall community structure

*Blautia* appears to be the most integral group for the gut of both ethnicities, since this genus was ranked as the most central taxon in both networks [68]. Subsequent Affinity Propagation [69] analysis revealed the sixth ranked node in the Dutch network, *Blautia massiliensis*, as the centroid of this large cluster.

*Coprococcus comes* was ranked as the second most central node in SAS and was unranked in the Dutch network. In the SAS, *C. comes* negatively correlated with two other taxa (*Anaerostipes hadrus* and *Clostridium sensu stricto 1*). In the Dutch, this bacterium positively correlated with eight other taxa (*Agathobacter, Bacteroides, Blautia massiliensis, Blautia, Anaerostipes hadrus, Dorea formicigenerans, Lachnospira,* and UCG-002) and negatively correlated with one other taxon (*Erysipelotrichaceae* UCG-003). *Oscillibacter* was another group that changed its direction of correlation with other bacteria between the two ethnicities, as it held a positive correlation with *Blautia massiliensis* in the Dutch and a negative correlation with the *Christensenellaceae* R7 group in the SAS.

*Haemophilus* and *Bifidobacterium* shared several positive edges with multiple other highly ranked nodes in the SAS network, and, in the Dutch, these genera were completely disconnected from the other nodes. In the SAS, *Haemophilus* shared positive edges with *Streptococcus* (ranked third) and *Anaerostipes hadrus* (ranked eleventh) while *Bifidobacterium* shared positive edges with *Blautia* (ranked first), *Blautia massiliensis* (ranked fourth), and *Agathobacter* (ranked 15th). A similar phenomenon occurred with *Bacteroides* and *Erysipelotrichaceae* UCG-003.

In the Dutch network, *Erysipelotrichaceae* UCG-003 (ranked fourth) was negatively correlated with eight other taxa (*Agathobacter, Bacteroides, Oscillibacter, Blautia, Dorea formicigenerans, Lachnospira*, and UCG-002), and positively correlated with one other taxon (*Marvinbryantia*). Also in the Dutch, *Bacteroides* (ranked third) held eight positive edges (*Agathobacter, Coprococcus comes*, UCG-002, *Anaerostipes hadrus, Lachnospira, Dorea formicigenerans, Blautia*, and *Blautia massiliensis)* and two negative edges (*Marvinbryantia* and *Erysipelotrichaceae* UCG-003). In the SAS, *Bacteroides* and *Erysipelotrichaceae* UCG-003 were not correlated with any other nodes.

## Discussion

The T2DM gut microbiome, in general, has been shown to be distinctly different from that of normoglycemic, insulin-sensitive individuals [15–17,22]. However, consensus on which taxa are enriched versus depleted as compared to healthy controls has yet to be reached [24]. Higher microbial production of SCFAs in the gut lumen is protective against T2DM [27,28]. The dynamics of SCFA-producing taxa have not been well-documented in minority populations because many microbiome studies are largely based on cohorts consisting of ethnic majority groups. It is important to understand how these taxa change with insulin resistance across all populations to develop effective and equitable microbiome-based therapeutics. In this study, the differential bacteria and ecological framework of the gut microbiota from two ethnic groups, the Dutch and South-Asian Surinamese (SAS), living in the same city (Amsterdam, the Netherlands) were analyzed. These two groups, both from the HELIUS cohort [39], were compared as the SAS had a much greater prevalence of T2DM. Microbiota compositions were generated via DADA2 (v1.16) [42]. Abundances were clustered using PCoA [45]. Metrics of alpha– and beta-diversity were computed. A paired t-test and two algorithms, DESeq2 [54,55] and LEfSe [53], were used to estimate differentially abundant taxa, or potential biomarkers. Lastly, bacterial co-occurrence networks [58] were constructed to examine the gut microbial ecology.

## Distinct SCFA-Producing Gut Microbial Milieus Between Ethnicities

The DESeq2 analysis (Table 3) identified several taxa that are capable of SCFA production to be in significantly higher abundance in the SAS gut. These included *Bifidobacterium*, *Anaerostipes*, and certain strains of *Haemophilus* [70–75]. *Clostridium sensu stricto 1* was detected as significantly more abundant in the Dutch, and it is unclear if this is a SCFA-producing taxon [76–78]. So, per the DESeq2 analysis, SCFA producers appeared to be in higher abundance in the SAS gut. Contrarily, the LEfSe biomarker analysis (Fig. 4) clearly discriminated the Dutch gut from that of the SAS by multiple SCFA-producers. These included *Butyricicoccus, Coprococcus comes, Lachnospira*, and *Odoribacter splanchnicus* [79–84]*. Alistipes putredinis* was also proposed as a taxon that could differentiate the Dutch gut, but this species is thought to produce SCFAs in much smaller quantities [85]. Other than *Haemophilus*, of which only certain strains produce SCFAs [75], no SCFA-producing taxa were proposed as biomarkers of the SAS gut microbiota by the LEfSe.

It is unclear from these opposing results if the overall abundances (Fig. 5) of SCFA-producing taxa truly vary between the two ethnicities. Various differential abundance methods are known to output different results [86]. Though, of the potential biomarkers identified, *Haemophilus parainfluenzae* ATCC 33392 (Table 4) would likely serve as the best discriminating bacterium between these two ethnicities because both differential abundance methods and a paired t-test between the RAs of the two groups agreed on its significantly higher abundance in the SAS gut. This was the only taxon on which all three tests of significance, i.e., the LEfSe, DESeq2, and paired t-test, reached consensus on differential abundance. In future work, it would be beneficial to add more methods, such as ALDEx2 [87] or ANCOM-BC [88], to identify with high certainty differential taxa between distinct microbial communities. While SCFA-producers may not be present in significantly different quantities between the Dutch and SAS, the Dutch gut certainly contained a more phylogenetically diverse SCFA-producing microbial milieu.

*Haemophilus parainfluenzae*, a known respiratory pathobiont, is also a gut pathobiont, as it plays a pro-inflammatory role in Crohn’s Disease [89,90]. A higher abundance of *Haemophilus parainfluenzae* has been significantly positively correlated with obesity and cardiometabolic disease by some [91], but negatively associated with similar entities by others [92,93]. To the best of our knowledge, *Haemophilus parainfluenzae* has not been clearly shown to produce SCFAs in a significant quantity. Our work indicates that this bacterium could be significantly associated, or even a pathobiont, in the guts of those with risk factors for metabolic disease, especially in ethnic minorities.

HbA1c, the most widely accepted marker of average blood glucose over time, has been shown to improve with complex fiber supplementation by upregulating microbial pathways for colonic SCFA fermentation [28,94]. SCFAs can potentiate glucose-stimulated insulin secretion, promote lipolysis, attenuate the T2DM-associated chronic inflammatory state, and modulate satiety [28,95]. *Bifidobacterium* and *Anaerostipes*, both SCFA-producers and previously negatively correlated with T2DM [23,25,28,96], were found to be significantly increased in the SAS [97]. Therefore, it is possible that T2DM-protective taxa may not play the same role in ethnic minority as compared to majority groups.

Whether certain microbes protect or predispose to T2DM is likely species– and/or strain-specific. Improved methods for easily identifying taxa at lower phylogenetic levels are necessary. *Bacteroides caccae* and *Lachnnospira eligens*, both SCFA-producers and previously negatively associated with T2DM [23,25,96], differentiated the Dutch gut per the LEfSe biomarker analysis [98]. However, at the genus level, *Bacteroides* was in significantly higher abundance in the SAS (Fig. 2 and 5), which further demonstrates that the relationship between taxon and host phenotype can only be inferred at the species– and/or strain-levels. The metabolically protective role of these species in previous works are likely robust findings. In general, the higher prevalence of T2DM seen in the SAS population in Amsterdam [41] may be associated with the loss of key SCFA producers.

## A More Complex Gut Microbiome Ecological Framework in the Ethnic Majority

Our PCoA analysis (Fig. 1) displayed that bacterial abundances and phylogeny were more similar across the Dutch samples as compared to the SAS. Sex was not observed to significantly impact gut microbiota because each PCoA cluster contained a roughly equal proportion of both sexes. This is consistent with previous work that has failed to demonstrate a clear relationship between sex and the gut microbiome [62]. Additionally, the overlap between the Dutch and SAS abundances may be explained by how long ago the SAS individuals had immigrated to the Netherlands, as over time they likely increasingly incorporated Dutch foods and were exposed to the Dutch environmental microbiome.

Bacterial co-occurrence, or social, networks of the Dutch and SAS gut microbiota were computed using RA data at the lowest possible taxonomic classification level (Fig. 6). Network analyses can help elucidate the ecological core microbiota [99], i.e., which taxa are key for community structure and, possibly, function. Compared to the SAS, the intra-taxonomic relationships in the Dutch gut were more interconnected, with many more positive correlational relationships. The Dutch network also contained a large cluster of SCFA-producing taxa (*Coprococcus comes, Lachnospira, Blautia,* and *Haemophilus*). This positively interconnected ecological framework correlates with the significantly higher average alpha diversity indices (Table 2) and more stable community composition and phylogenetic distribution across the Dutch samples (Fig. 1). Previous studies have associated diabetics with a less diverse gut microbiota [24,94].

Our networks support the notion that the role played, i.e., cooperation versus competition, by a certain taxon may change with ethnicity and is possibly a function of host glycemic phenotype. In the Dutch network, *Erysipelotrichaceae* UCG-003 was estimated to be the fourth most important taxon for the community structure and had several negative relationships with other taxa. However, in the SAS network, this taxon was unranked, and lacked any correlational relationships with other community members. *Erysieplotrichaceae* UCG-003 has previously been positively associated with insulin resistance and obesity [100,101]. *Erysieplotrichaceae* UCG-003 has also been proposed as a marker of healthy aging [102]. This taxon may be negatively correlated with others in the Dutch network because there is competition for a mutual resource. In the Dutch, the other likely beneficial taxa that *Erysieplotrichaceae* UCG-003 is correlated with may be better able to acquire that resource and use it to produce a metabolite that is beneficial for the host, such as a SCFA. This competition is perhaps lost in the SAS.

The Dutch, which are less afflicted with T2DM, may have a more interconnected and preserved gut microbial ecology because of unperturbed bacterial cross-feeding interactions. Cross-feeding is dependent on microbiota spatial organization, and is critical for SCFA production [27,28]. Several taxa that were either SCFA producers or negatively associated with T2DM were members of the large, interconnected Dutch network. *Bacteroides,* which is strongly negatively associated with T2DM [23,25,96], was completely disconnected and unranked in the SAS network, but was ranked as the third most integral taxon in the Dutch network. So, the metabolically beneficial role of this bacterium is likely a reproducible and accurate finding. *Oscillibacter,* ranked sixth in the Dutch and ninth in the SAS network, has been shown to be positively associated with T2DM [103,104]. *Bifidobacterium*, which has been reported to be protective against T2DM [14,23], had more positive relationships with other bacteria in the SAS group, but was unranked in both networks. So, the role of a particular taxon in relation to metabolic fitness may depend on host ethnicity.

The main limitation of our study was that we were unable to stratify the two ethnicities by diabetic status because we did not have access to their markers of hyperglycemia and insulin resistance, such as HbA1c and fasting plasma glucose. Although subjects were strictly age– and gender-matched, other factors, such as ethnically variable genetic predispositions [105,106], epigenetics [107,108], diet [61,109,110], and socioeconomic status [111,112], likely confounded our findings. The ethnic minorities of the HELIUS cohort were of lower socioeconomic status, which significantly impacts access to healthcare and health equity [41,107]. Differences in dietary and exercise patterns between the ethnicities were also observed in the HELIUS study [107]. Additionally, most of the SAS were first-generation immigrants, so they had a shorter length of residence in the Netherlands [39]. As the burden of T2DM is increasing across the globe, especially in developed countries, more work is needed to elucidate how the gut microbiome factors into the disease pathophysiology, so that additional anti-diabetic agents can be developed, such as pre– and probiotics.

## Statement and declarations

### Author contributions

Conceptualization: Eric I. Nayman, Brooke A. Schwartz, Michaela Polmann, Alayna C. Gumabong, and Kalai Mathee; Methodology: Eric I. Nayman, Brooke A. Schwartz, Max Nieuwdorp, and Trevor Cickovski; Data analysis: Eric I. Nayman, Brooke A. Schwartz, Michaela Polmann, Alayna C. Gumabong, Max Nieuwdorp, Trevor Cickovski, and Kalai Mathee; Writing and editing: Eric Nayman, Brooke A. Schwartz, Max Nieuwdorp, Trevor Cickovski, and Kalai Matheee; Supervision: Trevor Cickovski and Kalai Mathee. All authors have read and agreed to the published version of the manuscript.

### Ethics approval and consent to participate

The HELIUS study was approved by the medical ethics committee of the Amsterdam University Medical Centre, and all participants provided informed consent prior to enrollment in the study.

### Consent to publish

All authors consent for publication.

## List of abbreviations

ASV: Amplicon Sequence Variant
BP: base-pair
DESeq: Differential Expression Analysis for Sequence Count Data
GLP-1: Glucagon-Like Peptide 1
HELIUS: Healthy Life in an Urban Setting
PCoA: Principal Coordinate Analysis
RA: Relative Abundance
SAS: South-Asian Surinamese
SCFA: Short-Chain Fatty Acid
T2DM: Type Two Diabetes Mellitus

## Acknowledgements

The HELIUS study was conducted by the Amsterdam University Medical Centers, location AMC and the Public Health Service of Amsterdam. Both organizations provided core support for HELIUS. We are most grateful to the participants of the HELIUS study and the management team, research nurses, interviewers, research assistants, and other staff who have taken part in gathering the data of this study. We would like to thank the members of the FIU Bioinformatics Research Group for their valuable feedback throughout several discussions of this work.

## Data availability

<u>Data</u>: The HELIUS data are owned by the Amsterdam University Medical Centers, location AMC, in Amsterdam, the Netherlands. Any researcher can request the data by submitting a proposal to the HELIUS Executive Board, as outlined at http://www.heliusstudy.nl/en/researchers/collaboration, by email to. The HELIUS Executive Board will check proposals for compatibility with the general objectives, ethical approvals, and informed consent forms of the HELIUS study. There are no other restrictions to obtaining the data and all data requests will be processed in the same manner. The microbial genomic sequences from the HELIUS cohort, which were used for this study, are stored under protected access on the European Genome-Phenome Archive (https://ega-archive.org/datasets/EGAD00001004106).

Code/Software: Our entire downstream analysis through the open-source PluMA [1] initiative is available at http://biorg.cs.fiu.edu/pluma/pipelines.html.

## References

[1] Cickovski T, Narasimhan G. Constructing lightweight and flexible pipelines using Plugin-Based Microbiome Analysis (PluMA). Bioinformatics 2018;34:2881–8. 10.1093/bioinformatics/bty198.

[2] Berg G, Rybakova D, Fischer D, Cernava T, Vergès M-CC, Charles T, et al. Microbiome definition re-visited: old concepts and new challenges. Microbiome 2020;8:103. 10.1186/s40168-020-00875-0.

[3] Münger E, Montiel-Castro AJ, Langhans W, Pacheco-López G. Reciprocal Interactions Between Gut Microbiota and Host Social Behavior. Frontiers in Integrative Neuroscience 2018;12.

[4] Ogunrinola GA, Oyewale JO, Oshamika OO, Olasehinde GI. The Human Microbiome and Its Impacts on Health. International Journal of Microbiology 2020;2020:e8045646. 10.1155/2020/8045646.

[5] Hooks KB, O’Malley MA. Dysbiosis and Its Discontents. mBio 2017;8:e01492–17. 10.1128/mBio.01492-17.

[6] Foster JA, McVey Neufeld K-A. Gut–brain axis: how the microbiome influences anxiety and depression. Trends in Neurosciences 2013;36:305–12. 10.1016/j.tins.2013.01.005.

[7] Scher JU, Abramson SB. The microbiome and rheumatoid arthritis. Nat Rev Rheumatol 2011;7:569–78. 10.1038/nrrheum.2011.121.

[8] Wong SH, Yu J. Gut microbiota in colorectal cancer: mechanisms of action and clinical applications. Nat Rev Gastroenterol Hepatol 2019;16:690–704. 10.1038/s41575-019-0209-8.

[9] Khan I, Ullah N, Zha L, Bai Y, Khan A, Zhao T, et al. Alteration of Gut Microbiota in Inflammatory Bowel Disease (IBD): Cause or Consequence? IBD Treatment Targeting the Gut Microbiome. Pathogens 2019;8:126. 10.3390/pathogens8030126.

[10] Romano S, Savva GM, Bedarf JR, Charles IG, Hildebrand F, Narbad A. Meta-analysis of the Parkinson’s disease gut microbiome suggests alterations linked to intestinal inflammation. Npj Parkinsons Dis 2021;7:1–13. 10.1038/s41531-021-00156-z.

[11] Kang D-W, Adams JB, Gregory AC, Borody T, Chittick L, Fasano A, et al. Microbiota Transfer Therapy alters gut ecosystem and improves gastrointestinal and autism symptoms: an open-label study. Microbiome 2017;5:10. 10.1186/s40168-016-0225-7.

[12] Mathee K, Cickovski T, Deoraj A, Stollstorff M, Narasimhan G. The gut microbiome and neuropsychiatric disorders: implications for attention deficit hyperactivity disorder (ADHD). J Med Microbiol 2020;69:14–24. 10.1099/jmm.0.001112.

[13] Devaraj S, Hemarajata P, Versalovic J. The Human Gut Microbiome and Body Metabolism: Implications for Obesity and Diabetes. Clinical Chemistry 2013;59:617–28. 10.1373/clinchem.2012.187617.

[14] Bielka W, Przezak A, Pawlik A. The Role of the Gut Microbiota in the Pathogenesis of Diabetes. Int J Mol Sci 2022;23:480. 10.3390/ijms23010480.

[15] Vals-Delgado C, Alcala-Diaz JF, Molina-Abril H, Roncero-Ramos I, Caspers MPM, Schuren FHJ, et al. An altered microbiota pattern precedes Type 2 diabetes mellitus development: From the CORDIOPREV study. Journal of Advanced Research 2022;35:99–108. 10.1016/j.jare.2021.05.001.

[16] Fang Y, Zhang C, Shi H, Wei W, Shang J, Zheng R, et al. Characteristics of the Gut Microbiota and Metabolism in Patients With Latent Autoimmune Diabetes in Adults: A Case-Control Study. Diabetes Care 2021;44:2738–46. 10.2337/dc20-2975.

[17] Zhou W, Sailani MR, Contrepois K, Zhou Y, Ahadi S, Leopold SR, et al. Longitudinal multi-omics of host–microbe dynamics in prediabetes. Nature 2019;569:663–71. 10.1038/s41586-019-1236-x.

[18] National Diabetes Statistics Report | Diabetes | CDC 2022. https://www.cdc.gov/diabetes/data/statistics-report/index.html (accessed September 7, 2023).

[19] International Diabetes Federation (2019) IDF Diabetes Atlas: Ninth edition 2019. Available from https://diabetesatlas.org/upload/resources/material/20200302_133351_IDFATLAS9e-final-web.pdf. Accessed Sept 7th, 2023., n.d.

[20] Agyemang C, van der Linden EL, Bennet L. Type 2 diabetes burden among migrants in Europe: unravelling the causal pathways. Diabetologia 2021;64:2665–75. 10.1007/s00125-021-05586-1.

[21] DeFronzo RA, Ferrannini E, Groop L, Henry RR, Herman WH, Holst JJ, et al. Type 2 diabetes mellitus. Nat Rev Dis Primers 2015;1:15019. 10.1038/nrdp.2015.19.

[22] Cui J, Ramesh G, Wu M, Jensen ET, Crago O, Bertoni AG, et al. Butyrate-Producing Bacteria and Insulin Homeostasis: The Microbiome and Insulin Longitudinal Evaluation Study (MILES). Diabetes 2022;71:2438–46. 10.2337/db22-0168.

[23] Gurung M, Li Z, You H, Rodrigues R, Jump DB, Morgun A, et al. Role of gut microbiota in type 2 diabetes pathophysiology. eBioMedicine 2020;51. 10.1016/j.ebiom.2019.11.051.

[24] Letchumanan G, Abdullah N, Marlini M, Baharom N, Lawley B, Omar MR, et al. Gut Microbiota Composition in Prediabetes and Newly Diagnosed Type 2 Diabetes: A Systematic Review of Observational Studies. Frontiers in Cellular and Infection Microbiology 2022;12.

[25] Guo Z, Pan J, Zhu H, Chen Z-Y. Metabolites of Gut Microbiota and Possible Implication in Development of Diabetes Mellitus. J Agric Food Chem 2022;70:5945–60. 10.1021/acs.jafc.1c07851.

[26] Sanna S, van Zuydam NR, Mahajan A, Kurilshikov A, Vich Vila A, Võsa U, et al. Causal relationships among the gut microbiome, short-chain fatty acids and metabolic diseases. Nat Genet 2019;51:600–5. 10.1038/s41588-019-0350-x.

[27] Morrison DJ, Preston T. Formation of short chain fatty acids by the gut microbiota and their impact on human metabolism. Gut Microbes 2016;7:189–200. 10.1080/19490976.2015.1134082.

[28] Blaak E e., Canfora E e., Theis S, Frost G, Groen A k., Mithieux G, et al. Short chain fatty acids in human gut and metabolic health. Beneficial Microbes 2020;11:411–55. 10.3920/BM2020.0057.

[29] Kim CH. Microbiota or short-chain fatty acids: which regulates diabetes? Cell Mol Immunol 2018;15:88–91. 10.1038/cmi.2017.57.

[30] McNelis JC, Lee YS, Mayoral R, van der Kant R, Johnson AMF, Wollam J, et al. GPR43 Potentiates β-Cell Function in Obesity. Diabetes 2015;64:3203–17. 10.2337/db14-1938.

[31] Priyadarshini M, Villa SR, Fuller M, Wicksteed B, Mackay CR, Alquier T, et al. An Acetate-Specific GPCR, FFAR2, Regulates Insulin Secretion. Mol Endocrinol 2015;29:1055–66. 10.1210/me.2015-1007.

[32] Reddy MA, Chen Z, Park JT, Wang M, Lanting L, Zhang Q, et al. Regulation of inflammatory phenotype in macrophages by a diabetes-induced long noncoding RNA. Diabetes 2014;63:4249–61. 10.2337/db14-0298.

[33] Pingitore A, Chambers ES, Hill T, Maldonado IR, Liu B, Bewick G, et al. The diet-derived short chain fatty acid propionate improves beta-cell function in humans and stimulates insulin secretion from human islets in vitro. Diabetes Obes Metab 2017;19:257–65. 10.1111/dom.12811.

[34] Canfora EE, van der Beek CM, Jocken JWE, Goossens GH, Holst JJ, Olde Damink SWM, et al. Colonic infusions of short-chain fatty acid mixtures promote energy metabolism in overweight/obese men: a randomized crossover trial. Sci Rep 2017;7:2360. 10.1038/s41598-017-02546-x.

[35] van der Beek CM, Canfora EE, Lenaerts K, Troost FJ, Olde Damink SWM, Holst JJ, et al. Distal, not proximal, colonic acetate infusions promote fat oxidation and improve metabolic markers in overweight/obese men. Clinical Science 2016;130:2073–82. 10.1042/CS20160263.

[36] Freeland KR, Wolever TMS. Acute effects of intravenous and rectal acetate on glucagon-like peptide-1, peptide YY, ghrelin, adiponectin and tumour necrosis factor-alpha. Br J Nutr 2010;103:460–6. 10.1017/S0007114509991863.

[37] Louis P, Flint HJ. Formation of propionate and butyrate by the human colonic microbiota. Environ Microbiol 2017;19:29–41. 10.1111/1462-2920.13589.

[38] Gaulke CA, Sharpton TJ. The influence of ethnicity and geography on human gut microbiome composition. Nat Med 2018;24:1495–6. 10.1038/s41591-018-0210-8.

[39] Deschasaux M, Bouter KE, Prodan A, Levin E, Groen AK, Herrema H, et al. Depicting the composition of gut microbiota in a population with varied ethnic origins but shared geography. Nat Med 2018;24:1526–31. 10.1038/s41591-018-0160-1.

[40] Gacesa R, Kurilshikov A, Vich Vila A, Sinha T, Klaassen M a. Y, Bolte LA, et al. Environmental factors shaping the gut microbiome in a Dutch population. Nature 2022;604:732–9. 10.1038/s41586-022-04567-7.

[41] Snijder MB, Galenkamp H, Prins M, Derks EM, Peters RJG, Zwinderman AH, et al. Cohort profile: the Healthy Life in an Urban Setting (HELIUS) study in Amsterdam, The Netherlands. BMJ Open 2017;7. 10.1136/bmjopen-2017-017873.

[42] Callahan BJ, McMurdie PJ, Rosen MJ, Han AW, Johnson AJA, Holmes SP. DADA2: High-resolution sample inference from Illumina amplicon data. Nat Methods 2016;13:581–3. 10.1038/nmeth.3869.

[43] Quast C, Pruesse E, Yilmaz P, Gerken J, Schweer T, Yarza P, et al. The SILVA ribosomal RNA gene database project: improved data processing and web-based tools. Nucleic Acids Research 2013;41:D590–6. 10.1093/nar/gks1219.

[44] Neu AT, Allen EE, Roy K. Defining and quantifying the core microbiome: Challenges and prospects. Proceedings of the National Academy of Sciences 2021;118:e2104429118. 10.1073/pnas.2104429118.

[45] Pearson K. LIII. On lines and planes of closest fit to systems of points in space. The London, Edinburgh, and Dublin Philosophical Magazine and Journal of Science 1901;2:559–72. 10.1080/14786440109462720.

[46] Custer GF, Gans M, van Diepen LTA, Dini-Andreote F, Buerkle CA. Comparative Analysis of Core Microbiome Assignments: Implications for Ecological Synthesis. mSystems 2023;8:e01066–22. 10.1128/msystems.01066-22.

[47] Chao A. Nonparametric Estimation of the Number of Classes in a Population. Scandinavian Journal of Statistics 1984;11:265–70.

[48] Shannon CE. A mathematical theory of communication. The Bell System Technical Journal 1948;27:379–423. 10.1002/j.1538-7305.1948.tb01338.x.

[49] Fisher RA, Corbet AS, Williams CB. The Relation Between the Number of Species and the Number of Individuals in a Random Sample of an Animal Population. Journal of Animal Ecology 1943;12:42–58. 10.2307/1411.

[50] Simpson EH. Measurement of Diversity. Nature 1949;163:688–688. 10.1038/163688a0.

[51] Lozupone C, Knight R. UniFrac: a New Phylogenetic Method for Comparing Microbial Communities. Applied and Environmental Microbiology 2005;71:8228–35. 10.1128/AEM.71.12.8228-8235.2005.

[52] Lozupone C, Lladser ME, Knights D, Stombaugh J, Knight R. UniFrac: an effective distance metric for microbial community comparison. ISME J 2011;5:169–72. 10.1038/ismej.2010.133.

[53] Segata N, Izard J, Waldron L, Gevers D, Miropolsky L, Garrett WS, et al. Metagenomic biomarker discovery and explanation. Genome Biology 2011;12:R60. 10.1186/gb-2011-12-6-r60.

[54] Anders S, Huber W. Differential expression analysis for sequence count data. Genome Biology 2010;11:R106. 10.1186/gb-2010-11-10-r106.

[55] Love MI, Huber W, Anders S. Moderated estimation of fold change and dispersion for RNA-seq data with DESeq2. Genome Biology 2014;15:550. 10.1186/s13059-014-0550-8.

[56] Schoch CL, Ciufo S, Domrachev M, Hotton CL, Kannan S, Khovanskaya R, et al. NCBI Taxonomy: a comprehensive update on curation, resources and tools. Database (Oxford) 2020;2020:baaa062. 10.1093/database/baaa062.

[57] Faust K, Sathirapongsasuti JF, Izard J, Segata N, Gevers D, Raes J, et al. Microbial Co-occurrence Relationships in the Human Microbiome. PLOS Computational Biology 2012;8:e1002606. 10.1371/journal.pcbi.1002606.

[58] Fernandez M, Riveros JD, Campos M, Mathee K, Narasimhan G. Microbial “social networks.” BMC Genomics 2015;16:S6. 10.1186/1471-2164-16-S11-S6.

[59] Friedman J, Alm EJ. Inferring Correlation Networks from Genomic Survey Data. PLOS Computational Biology 2012;8:e1002687. 10.1371/journal.pcbi.1002687.

[60] Fruchterman TMJ, Reingold EM. Graph drawing by force-directed placement. Software: Practice and Experience 1991;21:1129–64. 10.1002/spe.4380211102.

[61] Nagpal R, Mainali R, Ahmadi S, Wang S, Singh R, Kavanagh K, et al. Gut microbiome and aging: Physiological and mechanistic insights. Nutr Healthy Aging 2018;4:267–85. 10.3233/NHA-170030.

[62] Kim YS, Unno T, Kim B-Y, Park M-S. Sex Differences in Gut Microbiota. World J Mens Health 2020;38:48–60. 10.5534/wjmh.190009.

[63] Snijder MB, Agyemang C, Peters RJ, Stronks K, Ujcic-Voortman JK, van Valkengoed IGM. Case Finding and Medical Treatment of Type 2 Diabetes among Different Ethnic Minority Groups: The HELIUS Study. J Diabetes Res 2017;2017:9896849. 10.1155/2017/9896849.

[64] Willis AD. Rarefaction, Alpha Diversity, and Statistics. Frontiers in Microbiology 2019;10.

[65] Janda JM, Abbott SL. 16S rRNA Gene Sequencing for Bacterial Identification in the Diagnostic Laboratory: Pluses, Perils, and Pitfalls. J Clin Microbiol 2007;45:2761–4. 10.1128/JCM.01228-07.

[66] Nearing JT, Douglas GM, Hayes MG, MacDonald J, Desai DK, Allward N, et al. Microbiome differential abundance methods produce different results across 38 datasets. Nat Commun 2022;13:342. 10.1038/s41467-022-28034-z.

[67] Cickovski T, Aguiar-Pulido V, Narasimhan G. MATria: a unified centrality algorithm. BMC Bioinformatics 2019;20:278. 10.1186/s12859-019-2820-7.

[68] Freeman LC. Centrality in social networks conceptual clarification. Social Networks 1978;1:215–39. 10.1016/0378-8733(78)90021-7.

[69] Frey BJ, Dueck D. Clustering by Passing Messages Between Data Points. Science 2007;315:972–6. 10.1126/science.1136800.

[70] Belenguer A, Duncan SH, Calder AG, Holtrop G, Louis P, Lobley GE, et al. Two Routes of Metabolic Cross-Feeding between Bifidobacterium adolescentis and Butyrate-Producing Anaerobes from the Human Gut. Appl Environ Microbiol 2006;72:3593–9. 10.1128/AEM.72.5.3593-3599.2006.

[71] Duncan SH, Louis P, Flint HJ. Lactate-utilizing bacteria, isolated from human feces, that produce butyrate as a major fermentation product. Appl Environ Microbiol 2004;70:5810–7. 10.1128/AEM.70.10.5810-5817.2004.

[72] Lee J-Y, Kang W, Shin N-R, Hyun D-W, Kim PS, Kim HS, et al. Anaerostipes hominis sp. nov., a novel butyrate-producing bacteria isolated from faeces of a patient with Crohn’s disease. Int J Syst Evol Microbiol 2021;71. 10.1099/ijsem.0.005129.

[73] Sato T, Matsumoto K, Okumura T, Yokoi W, Naito E, Yoshida Y, et al. Isolation of lactate-utilizing butyrate-producing bacteria from human feces and in vivo administration of Anaerostipes caccae strain L2 and galacto-oligosaccharides in a rat model. FEMS Microbiol Ecol 2008;66:528–36. 10.1111/j.1574-6941.2008.00528.x.

[74] Schwiertz A, Hold GL, Duncan SH, Gruhl B, Collins MD, Lawson PA, et al. Anaerostipes caccae gen. nov., sp. nov., a new saccharolytic, acetate-utilising, butyrate-producing bacterium from human faeces. Syst Appl Microbiol 2002;25:46–51. 10.1078/0723-2020-00096.

[75] López-López N, Euba B, Hill J, Dhouib R, Caballero L, Leiva J, et al. Haemophilus influenzae Glucose Catabolism Leading to Production of the Immunometabolite Acetate Has a Key Contribution to the Host Airway–Pathogen Interplay. ACS Infect Dis 2020;6:406–21. 10.1021/acsinfecdis.9b00359.

[76] Huart J, Leenders J, Taminiau B, Descy J, Saint-Remy A, Daube G, et al. Gut Microbiota and Fecal Levels of Short-Chain Fatty Acids Differ Upon 24-Hour Blood Pressure Levels in Men. Hypertension 2019;74:1005–13. 10.1161/HYPERTENSIONAHA.118.12588.

[77] Wang Y, Leong LEX, Keating RL, Kanno T, Abell GCJ, Mobegi FM, et al. Opportunistic bacteria confer the ability to ferment prebiotic starch in the adult cystic fibrosis gut. Gut Microbes 2018;10:367–81. 10.1080/19490976.2018.1534512.

[78] Hu C, Niu X, Chen S, Wen J, Bao M, Mohyuddin SG, et al. A Comprehensive Analysis of the Colonic Flora Diversity, Short Chain Fatty Acid Metabolism, Transcripts, and Biochemical Indexes in Heat-Stressed Pigs. Frontiers in Immunology 2021;12.

[79] Amiri P, Hosseini SA, Ghaffari S, Tutunchi H, Ghaffari S, Mosharkesh E, et al. Role of Butyrate, a Gut Microbiota Derived Metabolite, in Cardiovascular Diseases: A comprehensive narrative review. Frontiers in Pharmacology 2022;12.

[80] Eeckhaut V, Van Immerseel F, Teirlynck E, Pasmans F, Fievez V, Snauwaert C, et al. Butyricicoccus pullicaecorum gen. nov., sp. nov., an anaerobic, butyrate-producing bacterium isolated from the caecal content of a broiler chicken. Int J Syst Evol Microbiol 2008;58:2799–802. 10.1099/ijs.0.65730-0.

[81] Nogal A, Louca P, Zhang X, Wells PM, Steves CJ, Spector TD, et al. Circulating Levels of the Short-Chain Fatty Acid Acetate Mediate the Effect of the Gut Microbiome on Visceral Fat. Frontiers in Microbiology 2021;12.

[82] Abdugheni R, Wang W-Z, Wang Y-J, Du M-X, Liu F-L, Zhou N, et al. Metabolite profiling of human-originated Lachnospiraceae at the strain level. iMeta 2022;1:e58. 10.1002/imt2.58.

[83] Hiippala K, Barreto G, Burrello C, Diaz-Basabe A, Suutarinen M, Kainulainen V, et al. Novel Odoribacter splanchnicus Strain and Its Outer Membrane Vesicles Exert Immunoregulatory Effects in vitro. Frontiers in Microbiology 2020;11.

[84] Werner H, Rintelen G, Kunstek-Santos H. [A new butyric acid-producing bacteroides species: B. splanchnicus n. sp. (author’s transl)]. Zentralbl Bakteriol Orig A 1975;231:133–44.

[85] Parker BJ, Wearsch PA, Veloo ACM, Rodriguez-Palacios A. The Genus Alistipes: Gut Bacteria With Emerging Implications to Inflammation, Cancer, and Mental Health. Frontiers in Immunology 2020;11.

[86] Nearing JT, Douglas GM, Hayes MG, MacDonald J, Desai DK, Allward N, et al. Microbiome differential abundance methods produce different results across 38 datasets. Nat Commun 2022;13:342. 10.1038/s41467-022-28034-z.

[87] ANOVA-Like Differential Expression (ALDEx) Analysis for Mixed Population RNA-Seq | PLOS ONE n.d. https://journals.plos.org/plosone/article?id=10.1371/journal.pone.0067019 (accessed September 13, 2023).

[88] Lin H, Peddada SD. Analysis of compositions of microbiomes with bias correction. Nat Commun 2020;11:3514. 10.1038/s41467-020-17041-7.

[89] Sohn J, Li L, Zhang L, Genco RJ, Falkner KL, Tettelin H, et al. Periodontal disease is associated with increased gut colonization of pathogenic Haemophilus parainfluenzae in patients with Crohn’s disease. Cell Reports 2023;42:112120. 10.1016/j.celrep.2023.112120.

[90] Fitzgerald RS, Sanderson IR, Claesson MJ. Paediatric Inflammatory Bowel Disease and its Relationship with the Microbiome. Microb Ecol 2021;82:833–44. 10.1007/s00248-021-01697-9.

[91] de la Cuesta-Zuluaga J, Mueller NT, Álvarez-Quintero R, Velásquez-Mejía EP, Sierra JA, Corrales-Agudelo V, et al. Higher Fecal Short-Chain Fatty Acid Levels Are Associated with Gut Microbiome Dysbiosis, Obesity, Hypertension and Cardiometabolic Disease Risk Factors. Nutrients 2018;11:51. 10.3390/nu11010051.

[92] Chen J, Qin Q, Yan S, Yang Y, Yan H, Li T, et al. Gut Microbiome Alterations in Patients With Carotid Atherosclerosis. Frontiers in Cardiovascular Medicine 2021;8.

[93] Palmnäs-Bédard MSA, Costabile G, Vetrani C, Åberg S, Hjalmarsson Y, Dicksved J, et al. The human gut microbiota and glucose metabolism: a scoping review of key bacteria and the potential role of SCFAs. Am J Clin Nutr 2022;116:862–74. 10.1093/ajcn/nqac217.

[94] Makki K, Deehan EC, Walter J, Bäckhed F. The Impact of Dietary Fiber on Gut Microbiota in Host Health and Disease. Cell Host Microbe 2018;23:705–15. 10.1016/j.chom.2018.05.012.

[95] Roelofsen H, Priebe MG, Vonk RJ. The interaction of short-chain fatty acids with adipose tissue: relevance for prevention of type 2 diabetes. Benef Microbes 2010;1:433–7. 10.3920/BM2010.0028.

[96] Iatcu CO, Steen A, Covasa M. Gut Microbiota and Complications of Type-2 Diabetes. Nutrients 2022;14:166. 10.3390/nu14010166.

[97] Zeevi D, Korem T, Godneva A, Bar N, Kurilshikov A, Lotan-Pompan M, et al. Structural variation in the gut microbiome associates with host health. Nature 2019;568:43–8. 10.1038/s41586-019-1065-y.

[98] Maskarinec G, Raquinio P, Kristal BS, Setiawan VW, Wilkens LR, Franke AA, et al. The gut microbiome and type 2 diabetes status in the Multiethnic Cohort. PLoS One 2021;16:e0250855. 10.1371/journal.pone.0250855.

[99] Risely A. Applying the core microbiome to understand host–microbe systems. Journal of Animal Ecology 2020;89:1549–58. 10.1111/1365-2656.13229.

[100] Atzeni A, Bastiaanssen TFS, Cryan JF, Tinahones FJ, Vioque J, Corella D, et al. Taxonomic and Functional Fecal Microbiota Signatures Associated With Insulin Resistance in Non-Diabetic Subjects With Overweight/Obesity Within the Frame of the PREDIMED-Plus Study. Front Endocrinol (Lausanne) 2022;13:804455. 10.3389/fendo.2022.804455.

[101] Li Y, Kang Y, Du Y, Chen M, Guo L, Huang X, et al. Effects of Konjaku Flour on the Gut Microbiota of Obese Patients. Frontiers in Cellular and Infection Microbiology 2022;12.

[102] Singh H, Torralba MG, Moncera KJ, DiLello L, Petrini J, Nelson KE, et al. Gastro-intestinal and oral microbiome signatures associated with healthy aging. GeroScience 2019;41:907–21. 10.1007/s11357-019-00098-8.

[103] Wu X, Park S. Fecal Bacterial Community and Metagenome Function in Asians with Type 2 Diabetes, According to Enterotypes. Biomedicines 2022;10:2998. 10.3390/biomedicines10112998.

[104] Thingholm LB, Rühlemann MC, Koch M, Fuqua B, Laucke G, Boehm R, et al. Obese Individuals with and without Type 2 Diabetes Show Different Gut Microbial Functional Capacity and Composition. Cell Host Microbe 2019;26:252–264.e10. 10.1016/j.chom.2019.07.004.

[105] Langenberg C, Lotta LA. Genomic insights into the causes of type 2 diabetes. Lancet 2018;391:2463–74. 10.1016/S0140-6736(18)31132-2.

[106] Sirdah MM, Reading NS. Genetic predisposition in type 2 diabetes: A promising approach toward a personalized management of diabetes. Clin Genet 2020;98:525–47. 10.1111/cge.13772.

[107] Stronks K, Snijder MB, Peters RJ, Prins M, Schene AH, Zwinderman AH. Unravelling the impact of ethnicity on health in Europe: the HELIUS study. BMC Public Health 2013;13:402. 10.1186/1471-2458-13-402.

[108] Shojima N, Yamauchi T. Progress in genetics of type 2 diabetes and diabetic complications. J Diabetes Investig 2023;14:503–15. 10.1111/jdi.13970.

[109] Stephenson EJ, Smiles W, Hawley JA. The relationship between exercise,nutrition and type 2 diabetes. Med Sport Sci 2014;60:1–10. 10.1159/000357331.

[110] Delpino FM, Figueiredo LM, Bielemann RM, da Silva BGC, Dos Santos FS, Mintem GC, et al. Ultra-processed food and risk of type 2 diabetes: a systematic review and meta-analysis of longitudinal studies. Int J Epidemiol 2022;51:1120–41. 10.1093/ije/dyab247.

[111] Canedo JR, Miller ST, Schlundt D, Fadden MK, Sanderson M. Racial/Ethnic Disparities in Diabetes Quality of Care: the Role of Healthcare Access and Socioeconomic Status. J Racial Ethn Health Disparities 2018;5:7–14. 10.1007/s40615-016-0335-8.

[112] Espelt A, Arriola L, Borrell C, Larrañaga I, Sandín M, Escolar-Pujolar A. Socioeconomic position and type 2 diabetes mellitus in Europe 1999-2009: a panorama of inequalities. Curr Diabetes Rev 2011;7:148–58. 10.2174/157339911795843131.

